# Tactile enrichment drives emergence of functional columns and improves sensory coding in L2/3 of mouse S1

**DOI:** 10.1101/447508

**Authors:** Amy M. LeMessurier, Daniel E. Feldman

## Abstract

Sensory maps in layer (L) 2/3 of rodent cortex lack precise functional column boundaries, and instead exhibit locally heterogeneous tuning superimposed on smooth global topography. Could this organization be a byproduct of impoverished experience in laboratory housing? We compared whisker map somatotopy in L2/3 and L4 excitatory cells of somatosensory (S1) cortex in normally housed vs. tactile-enriched mice, using GCaMP6s imaging. Normally housed mice had a dispersed, salt-and-pepper whisker map in L2/3, but L4 was more topographically precise. Enrichment (P21 to P46-71) sharpened whisker tuning and decreased, but did not abolish, local tuning heterogeneity. In L2/3, enrichment strengthened and sharpened whisker point representations, and created functional boundaries of tuning similarity and noise correlations at column edges. Thus, tactile experience drives emergence of functional columnar topography in S1, and reduces salt-and-pepper tuning heterogeneity. These changes predict improved single-trial population coding of whisker deflections within each column.

## Introduction

Sensory maps are a defining feature of cortical sensory areas, and may enable efficient local network computations (Chklovskii and Koulakov, 2004). In rodents, maps in L2/3 of primary sensory cortex exhibit strong tuning heterogeneity between nearby neurons, superimposed on smooth global topography (Andermann et al., 2013; Bonin et al., 2011; Kanold et al., 2014; Ohki et al., 2005). In these salt-and-pepper maps, discrete functional column boundaries are lacking and the neural ensemble activated by a single sensory feature is spatially dispersed, though synaptically linked (Harris and Mrsic-Flogel, 2013). The origins and function of such dispersed maps are unclear. L2/3 maps are also robustly plastic to sensory experience (Buonomano and Merzenich, 1998; Feldman and Brecht, 2005; LeMessurier and Feldman, 2018). Appropriate sensory experience is often required for development of normal map topography (Katz and Shatz, 1996), raising the question of whether dispersed, salt-and-pepper maps are a natural feature of L2/3, or may reflect incomplete activity-dependent development due to impoverished sensory experience in rodent laboratory housing.

Sensory experience may be particularly relevant for development of whisker map somatotopy in rodent S1. Whiskers are tactile sensors represented in S1 by a topographic array of cortical columns, each centered on a L4 barrel. In the classical whisker map, inferred from single-unit recordings, nearly all neurons are tuned for their column’s anatomically corresponding whisker (CW). However, recent studies using cellular-resolution population calcium imaging in L2/3 reveal intermixed tuning in each column, with neurons tuned heterogeneously for either the CW or a surround whisker (SW). While average tuning within each column is topographically correct, deflection of one whisker activates a dispersed set of L2/3 neurons spread across multiple columns without discrete functional column borders (Clancy et al., 2015; Sato et al., 2007). This globally smooth but locally scattered representation resembles salt-and-pepper maps of orientation tuning and retinotopy in rodent V1 (Bonin et al., 2011; Ohki et al., 2005) and frequency tuning in rodent A1 (Kanold et al., 2014).

These dispersed L2/3 maps were discovered in rodents reared in standard laboratory housing, which is a relatively deprived sensory environment. Environmental enrichment increases the amount, complexity and salience of sensory stimuli, and promotes cortical development (Diamond et al., 1976). Enrichment sharpens single-unit receptive fields and average map topography in forepaw S1 and A1 of young rodents (Cai et al., 2009; Coq and Xerri, 1998; Engineer et al., 2005; Jakkamsetti et al., 2012), and in whisker S1 of adults (Polley et al., 2004). But how enrichment affects cellular-resolution maps is unknown. If the dispersed, salt-and-pepper whisker map in L2/3 is a byproduct of inadequate sensory experience, tactile enrichment should increase columnar structure and refine map topography at the cellular scale.

To test this question, we raised young adult mice from P21 for 25-50 days in either standard laboratory housing with a single littermate, or in an enriched environment with tactile toys and an additional littermate, which will further enhance whisker contact as part of social interactions (Rao et al., 2014). We then characterized whisker tuning, responsiveness, and map organization in L2/3 or L4 excitatory cells using 2-photon imaging of GCaMP6s expressed conditionally using layer-specific Cre lines. Results confirmed the dispersed, weakly columnar whisker map in L2/3 of normally housed (NH) mice. Enrichment (EN) strongly increased columnar organization and reduced, but did not eliminate, salt-and-pepper organization. These effects occurred mostly in L2/3, not L4. Thus, enrichment greatly enhances columnar organization in L2/3 of S1, so that cellular-scale functional topography better matches the anatomical columnar map.

## RESULTS

### Whisker map topography in L2/3 and L4 in normally housed mice

We measured whisker receptive fields and maps among L2/3 pyramidal (PYR) cells and L4 excitatory cells, by injecting Cre-dependent GCaMP6s viral vectors into S1 of L2/3 PYR-specific (Drd3-Cre) (Gong et al., 2007) or L4 excitatory-specific (Scnn1a-Tg3-Cre) mice (Adesnik et al., 2012; Madisen et al., 2010). After allowing several weeks for expression, we used resonant-scanned 2-photon calcium imaging to measure whisker-evoked neural activity through a chronic cranial window in anesthetized mice. L2/3 imaging fields were ~400 x 400 μ.m and contained ~50-100 expressing neurons. Smaller imaging fields were used in L4 (~300 x 300 μm) to compensate signal-to-noise ratio for imaging depth. Imaging was performed in D2 and adjacent whisker columns. We measured ΔF/F responses to deflection of 9 whiskers in a 3 x 3 grid centered on D2, using brief 5-pulse deflection trains, plus no-stimulation ‘blank’ trials **(Fig. 1A-C**). 50-70 repetitions of each whisker stimulus were randomly interleaved. Regions of interest (ROI) corresponding to GCaMP6-expressing somata were considered significantly whisker-responsive if the distribution of whisker-evoked ΔF/F (in a 1-sec window after whisker deflection) was greater than blank trials (permutation test, α = 0.05 corrected for 9 whiskers, see Methods). Analyses were restricted to whisker-responsive ROIs. Each cell’s receptive field was quantified from median whisker-evoked ΔF/F, and its best whisker (BW) was defined as that evoking the strongest median response. Three example cells from one imaging field are shown in **Fig. 1D**. Imaged neurons were localized post-hoc relative to L4 barrel column centers and boundaries using histological staining and alignment by surface blood vessels (**Fig. 1E**, see Methods). We confirmed that GCaMP6s-based receptive fields were similar to spiking-based receptive fields by performing simultaneous juxtacellular spike recording and GCaMP6s imaging in 11 L2/3 neurons. Receptive fields assessed by spike counts (0-100 ms after each whisker deflection) were highly similar to those based on ΔF/F for 10/11 neurons, and yielded the same BW for 8/11 cells (**Fig. 1F**).

**Figure 1.**
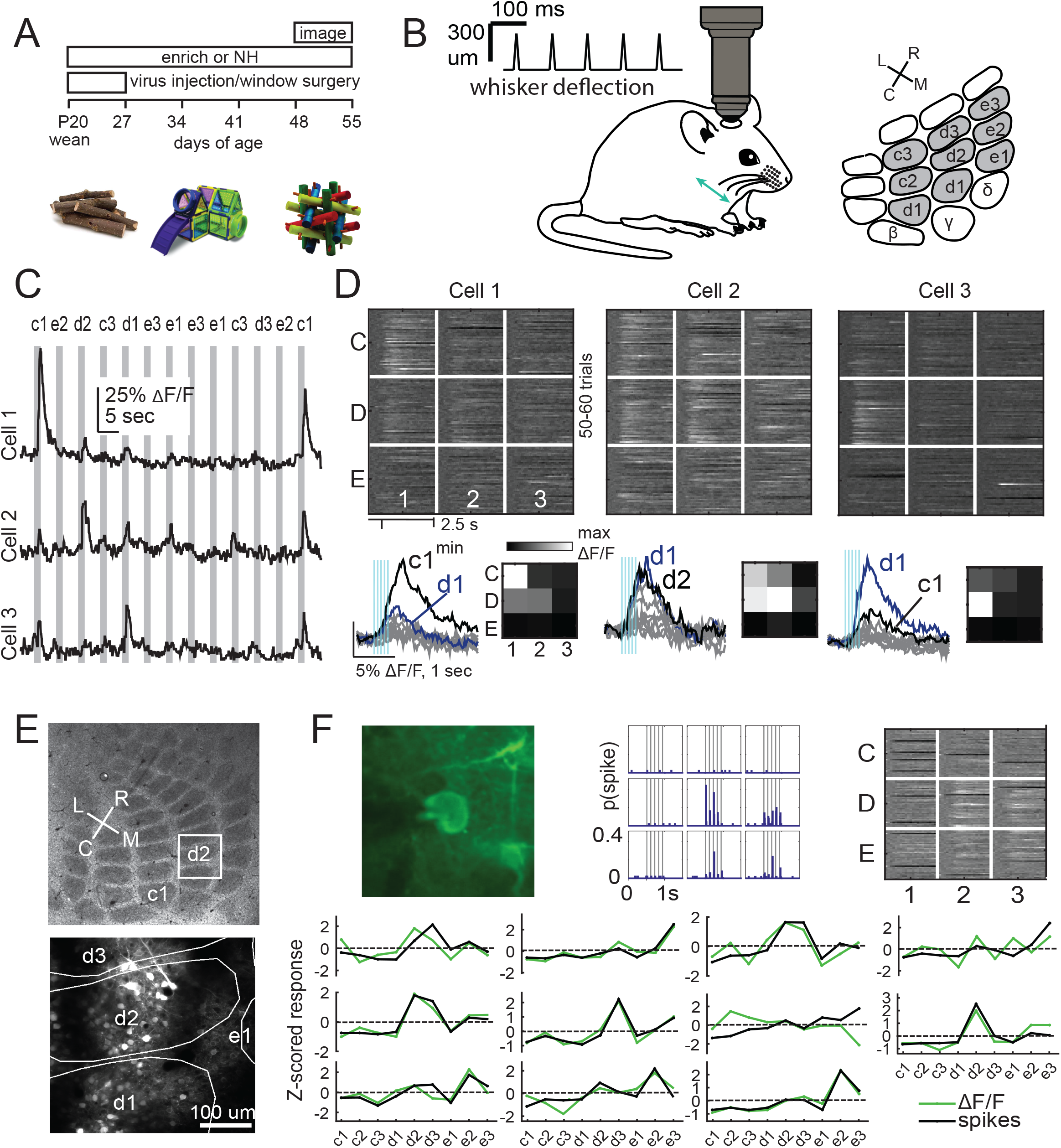
2-photon calcium imaging of whisker responses and receptive fields in S1. **(A)** Experimental time line and example enrichment objects. **(B)** Schematic of whisker stimulus train and barrel map showing the 3×3 array of stimulated whiskers in gray. **(C)** Example ΔF/F traces from 3 ROIs imaged simultaneously in L2/3 of one column. Grey bars are deflections of each indicated whisker. **(D)** Whisker receptive fields for the 3 cells in (C). Top, ΔF/F traces (shown as grayscale after baseline subtraction) for each trial for the 9 whiskers (C1 to E3), aligned to stimulus onset (St). Bottom left, Mean ΔF/F trace for each whisker. Vertical bars are whisker deflections. The 2 strongest whiskers are shown in black and blue, respectively. Bottom right, Median ΔF/F averaged over 1-sec response window. **(E)** Localization of one L2/3 imaging field (white box) to barrels recovered in a cytochrome oxidase-stained section through L4. Bottom, projection of barrel boundaries onto this imaging field. **(F)** Similarity of receptive fields measured by simultaneous GCaMP6s imaging and loose-seal cell-attached spike recording. Top row: One example L2/3 neuron shown during measurement, with its imaging receptive field (plotted as in D), and its spiking receptive field shown as a peristimulus time histogram (PSTH) for each whisker. Bottom row: Average imaging (red) and spiking (black) receptive fields for each neuron (n=11).

In L2/3, we quantified salt-and-pepper organization selectively among PYR cells, which has not been done previously (Clancy et al., 2015; Sato et al., 2007), by imaging in Drd3-Cre mice. In normally housed (NH) mice (age: P56 ± 11), 47.6% of L2/3 ROIs were responsive to at least one whisker. BW tuning was heterogeneous in individual columns. To quantify this, we calculated the fraction of cells tuned to a given whisker within spatial bins relative to the center of that whisker’s anatomical column. In L2/3, 49.5% (50/101) of cells located in the central bin of a column (0-91 μm) were tuned for that column’s whisker (i.e., the CW was the cell’s BW). This fraction fell off gradually with cortical distance to 26.3% at 182-273 μm and 2.6% at 455-546 μm (**Fig. 2A-B**). This distribution is broad compared to mean anatomical column radius (147 ± 11 μm). Because of statistical uncertainty in identifying the BW from the limited stimulus repetitions, we also identified all whiskers that evoked a mean ΔF/F that was statistically indistinguishable from the BW (see Methods). These were termed equal best whiskers (EBWs). In L2/3, 62.4% (63/101) of cells at the center of a column had the CW as an EBW. This fell off gradually to 48.6% at 182-273 μm and 14.1% (77/545 cells) at 455-546 μm from column center (**Fig. 2C-D**). Thus, in NH mice L2/3 PYR cells tuned for a given whisker were organized in a dispersed, salt-and-pepper fashion, consistent with prior results (Sato et al., 2007; Clancy et al., 2015).

**Figure 2.**
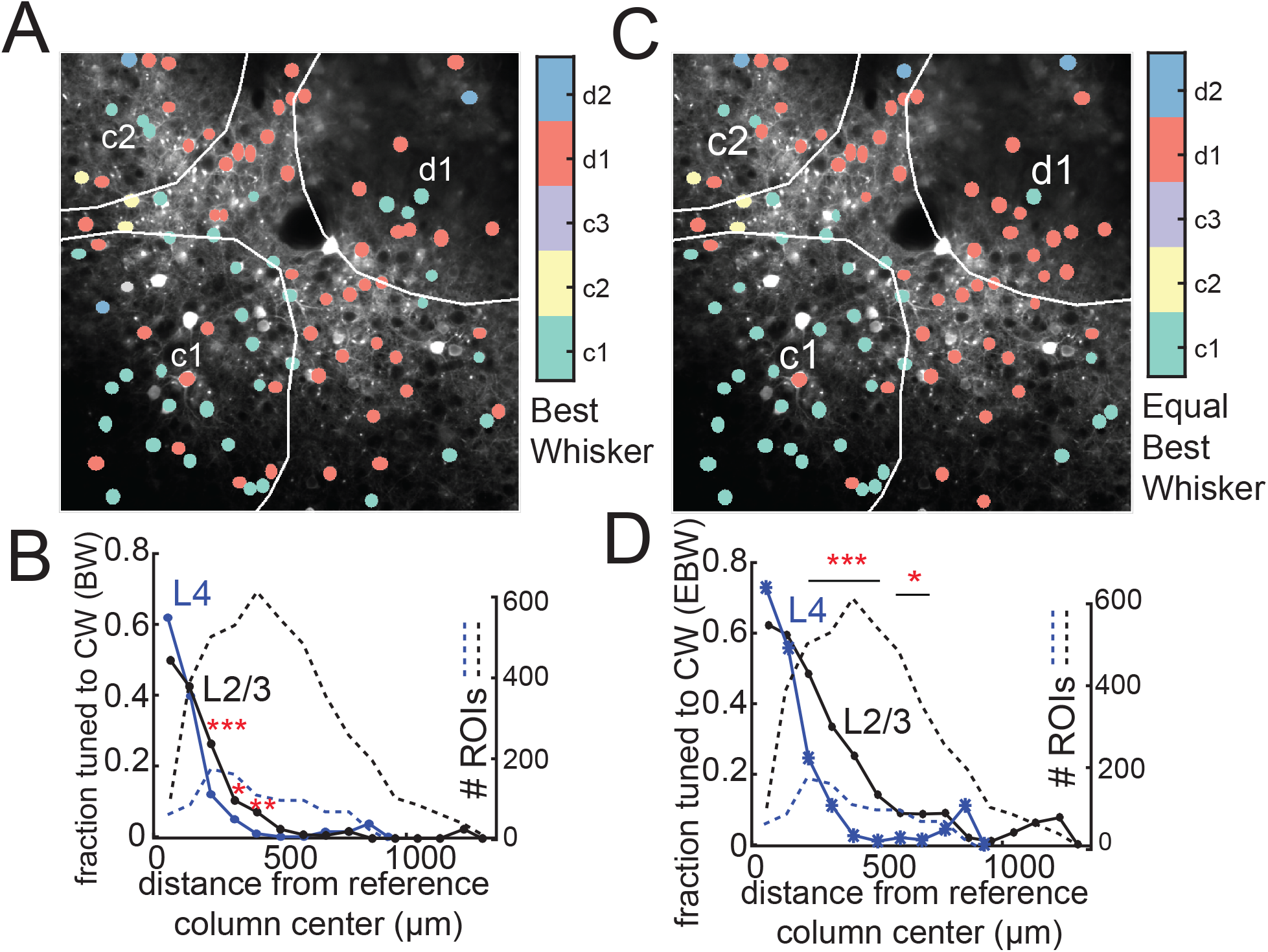
Salt-and-pepper tuning organization among L2/3 and L4 excitatory cells in normally housed mice. **(A)** Example L2/3 imaging field with ROIs color-coded by the whisker that evoked that largest absolute response. **(B)** Fraction of responsive excitatory ROIs that were tuned to a reference whisker (defined as evoking the absolute largest response) in spatial bins of distance from that whisker’s anatomical column center. L2/3, black. L4, blue. Dashed lines, number of ROIs in each bin. **(C)** Same imaging field as (A), with neurons inside each column re-colored to the columnar whisker if it was either the cell’s absolute best whisker, or evoked a response that was statistically indistinguishable from the absolute best whisker. These neurons are ‘equivalently tuned’ to the CW, meaning they are either tuned or co-tuned for the CW. **(D)** Fraction of L2/3 PYR ROIs equivalently tuned to a reference whisker as a function of distance from the reference whisker column center. Asterisks show significant differences between L4 and L2/3 by Fisher’ Exact Test, computed separately in each spatial bin. * p<0.05, ** p<0.01, *** p<0.001.

In L4, the column-specific termination of thalamic axons and prior physiology suggests a more precise whisker map (Andrew Hires et al., 2015; Simons, 1978). To test this, we imaged L4 excitatory neurons in Scnn1a-Cre mice. In NH mice, 60.3% of L4 ROIs were whisker-responsive. At the center of a barrel (0-91 um), 61.9% of cells (39/63) were tuned for the CW, and 73% (46/63) had the CW as an EBW. This precision is not different from L2/3 (p=0.15, Fisher’s Exact Test). But tuning fell off ~30% more sharply with distance in L4 than in L2/3, both for absolute BW and for EBW (**Fig. 2B & D**). Thus, at 182-273 μm from the column center, just 12% of L4 cells were tuned to CW (p=0.0001 vs. L2/3, Fisher’s exact test), and just 24.6% had the CW as an EBW (p=0, Fisher’s Exact Test). Thus, L4 exhibits more spatial map precision than L2/3.

### Enrichment decreases tuning heterogeneity in the salt-and-pepper map

To study effects of enrichment, littermates were separated into enriched (EN) and NH cohorts at weaning (P21). EN mice were housed with 2-3 littermates with tactile toys of varying material, textures, and geometry that were exchanged every 2-3 days to encourage exploration (**Fig. 1A**). NH mice were housed with a single littermate and standard wood chip bedding and nesting material. Imaging was performed after 25-50 days of EN or NH experience.

Enrichment decreased local tuning heterogeneity in each column in L2/3. In EN mice, 80.0% (168/210) of cells at column centers (0-91 μ.m) had the CW as an EBW, which is greater than NH mice (p=0.0013, Fisher’s Exact Test). While tuning precision was higher for several bins near the column center, it was not affected >172 μm from the column center (**Fig. 3A**). When all cells located within column boundaries are considered together (irrespective of distance to center), 77.1% (468/607) of whisker-responsive cells in EN mice had the CW as an EBW, compared to 65.1% (224/344) in NH (p=0.00008). Thus, enrichment increased L2/3 map precision, but did not abolish salt-and-pepper tuning. We define the tuning ensemble for one whisker as the set of cells with equivalent best tuning to that whisker. In NH mice, the tuning ensemble in L2/3 was dispersed across columns. EN increased the density of the tuning ensemble at its center (within the reference whisker’s home column), but did not alter its overall spread (**Fig. 3C**).

**Figure 3.**
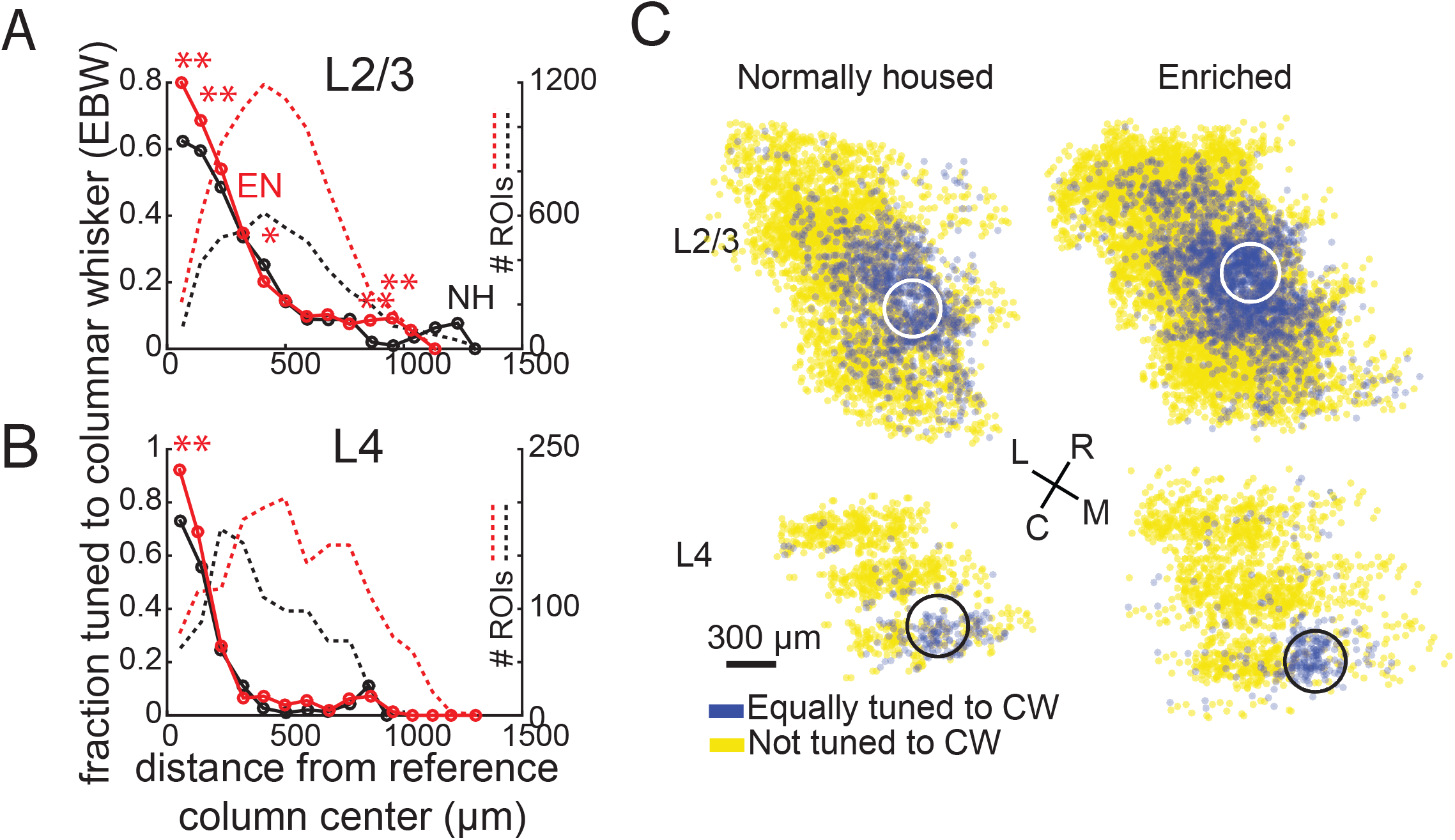
Effect of enrichment on somatotopic tuning precision. **(A)** Fraction of L2/3 PYR ROIs (top) or L4 excitatory ROIs (bottom) that were equivalently tuned to a reference whisker in EN and NH mice. Asterisks show significant differences between EN and NH by Fisher’ Exact Test, computed separately in each spatial bin. * p<0.05, ** p<0.01, *** p<0.001. **(B)** Spatial map of ROIs equivalently tuned for a reference whisker, relative to the center of that whisker’s column Data are combined across all imaging fields after spatial alignment. White or black circle is the reference column (shown as an average barrel diameter around the column center). ROIs are layered and semi-transparent, so that density of CW-tuned (blue) and non-CW-tuned (yellow) ROIs can be observed. Enrichment increases the density of ROIs equivalently tuned to a reference whisker in and just around its column.

Enrichment also decreased tuning heterogeneity in L4. In EN mice, 92.2% (71/77) of cells at column centers (0-91 μm) had the CW as an EBW, which is greater than the 73% in NH mice (p=0.0028). This effect was confined to this one bin at the column center (**Fig. 3B**). Considering all cells within barrel boundaries, 89.3% (133/149) of cells in EN mice had the CW as an EBW, compared to 75.6% (65/86) in NH (p=0.0086). Like in L2/3, this increased the density of L4 tuning ensembles at their centers (**Fig. 3C**). Thus, despite the expectation that L4 is less plastic than L2/3 (Feldman and Brecht, 2005; Glazewski and Fox, 1996), enrichment also increased tuning precision in L4.

### Enrichment sharpens whisker receptive fields

Enrichment sharpened L2/3 whisker receptive fields. Whisker responses in each cell were normalized to baseline (blank trial) activity, rank-ordered to quantify tuning sharpness independent of somatotopic shape or precision, and then averaged across cells. In L2/3, EN cells had greater responses to the strongest 3 whiskers, and weaker responses to the weakest 4 whiskers, than NH cells (**Fig. 4A**). EN specifically increased both CW response and the best 2 SW responses (**Fig. 4B**) In addition to these changes in tuning shape, EN also decreased spontaneous (blank trial) activity relative to NH (**Fig. 4C**). To test whether receptive field sharpening was evident in raw response amplitude (without normalization to spontaneous activity), we compared within each L2/3 cell the mean evoked ΔF/F for the 4 weakest whiskers vs. the 4 strongest whiskers. The slope of this relationship across cells was higher in EN than NH, indicating sharper tuning for raw evoked responses (NH: R^2^=0.12, slope = 0.67; EN: R^2^=0.28, slope= 1.24; p=2.14*10^−9^, t-test for difference in slopes). Sharpening of tuning was more prominent for L2/3 cells located over L4 barrels than for those located over L4 septa (**Fig. 4E**). Because a low fraction of imaged cells were located over septa (**Fig. 4F**), sharpening of tuning for barrel-related cells is likely responsible for the sharpening in the overall population.

**Figure 4.**
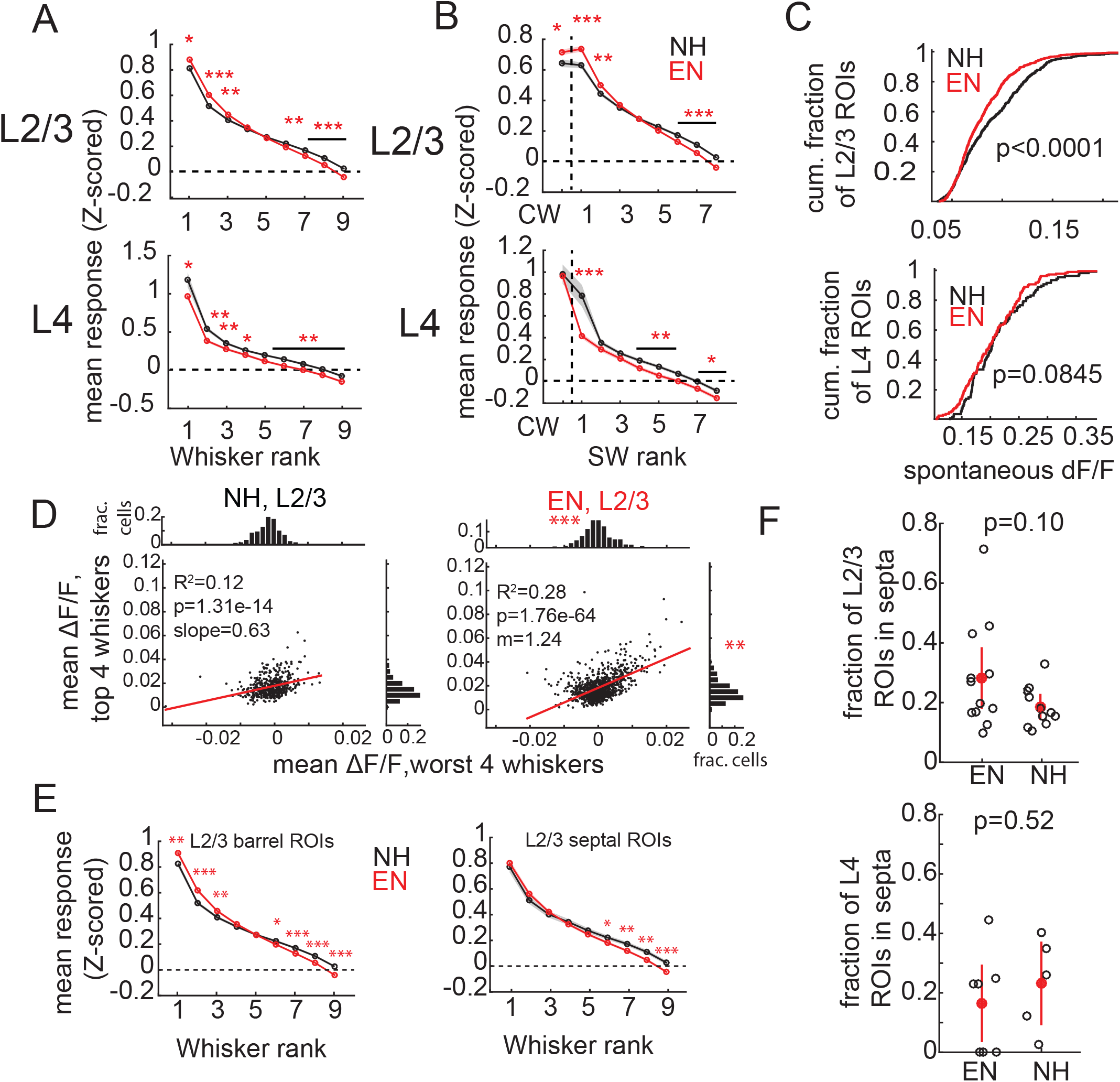
Enrichment sharpens whisker tuning curves. **(A)** Mean receptive field for all responsive ROIs, calculated after ranking whiskers from strongest to weakest in each cell and normalizing median responses to spontaneous activity. Asterisks, significant difference between EN and NH, computed separately by permutation test for each whisker rank. * p<0.05, ** p<0.01, *** p<0.001. **(B),** Mean receptive fields separated into CW and ranked SW whiskers. **(C)** Effect of enrichment on spontaneous activity, defined as median ΔF/F during blank trials. P-values computed by permutation test. **(D)** Correlation of mean ΔF/F evoked by 4 worst-ranked whiskers to 4 best-ranked whiskers within each cell. Each dot on scatter plots indicates one ROI. Red lines indicate linear regression fit. Histograms indicate distributions of raw evoked response magnitude. Asterisks denote difference between EN and NH distributions from permutation test. Note: in EN, one point at (0.03, 0.17) is omitted from scatter plot due to scale. **(E)** Effect of enrichment on average ranked whisker tuning curves for barrel- and septal-related ROIs in L2/3. (**F)** Fraction of whisker-responsive ROIs located in septa (L4) or over septa (L2/3) septa. Each circle represents one imaging field. Red is population mean and SEM. P-values computed from 2-sample t-test.

EN also sharpened L4 whisker receptive fields, but by a different mechanism. EN cells had reduced responses to most whiskers in rank-ordered receptive fields, except for CW responses, which were preserved (**Fig. 4A-B**). Spontaneous activity was unaffected (**Fig. 4C**). Thus, EN sharpened L4 receptive fields around the CW by weakening all but CW responses, on average. These results differ from adult enrichment in rats, which shrinks whisker receptive fields in L2/3, but not L4 (Polley et al., 2004).

### Enrichment sharpens the point representation of a single whisker in L2/3, but not L4

To assess whisker map topography we measured the point representation, i.e. the spatial profile of activity in S1 evoked by deflection of a single whisker. We quantified the point representation of a reference whisker as evoked ΔF/F relative to baseline, averaged across all whisker-responsive cells, as a function of distance from that whisker’s column center. In L2/3 of NH mice, the point representation was centered on the reference whisker’s anatomical column, and fell off gradually over several columns’ distance. EN mice showed an elevated mean response at the reference column center (0-91 μm: NH: 0.36 ± 0.03, EN: 0.47 ± 0.02) and for all spatial bins up to 364 μm (p<0.0002), which includes immediate neighboring columns. The more distant tail of the point representation was unchanged (**Fig. 5A**). These effects were also apparent in 2D maps in which ROIs were spatially binned along axes aligned to the reference column center (**Fig. 5B**). This is consistent with EN strengthening of both CW responses and the strongest SW responses in L2/3 receptive fields (**Fig. 4**).

**Figure 5.**
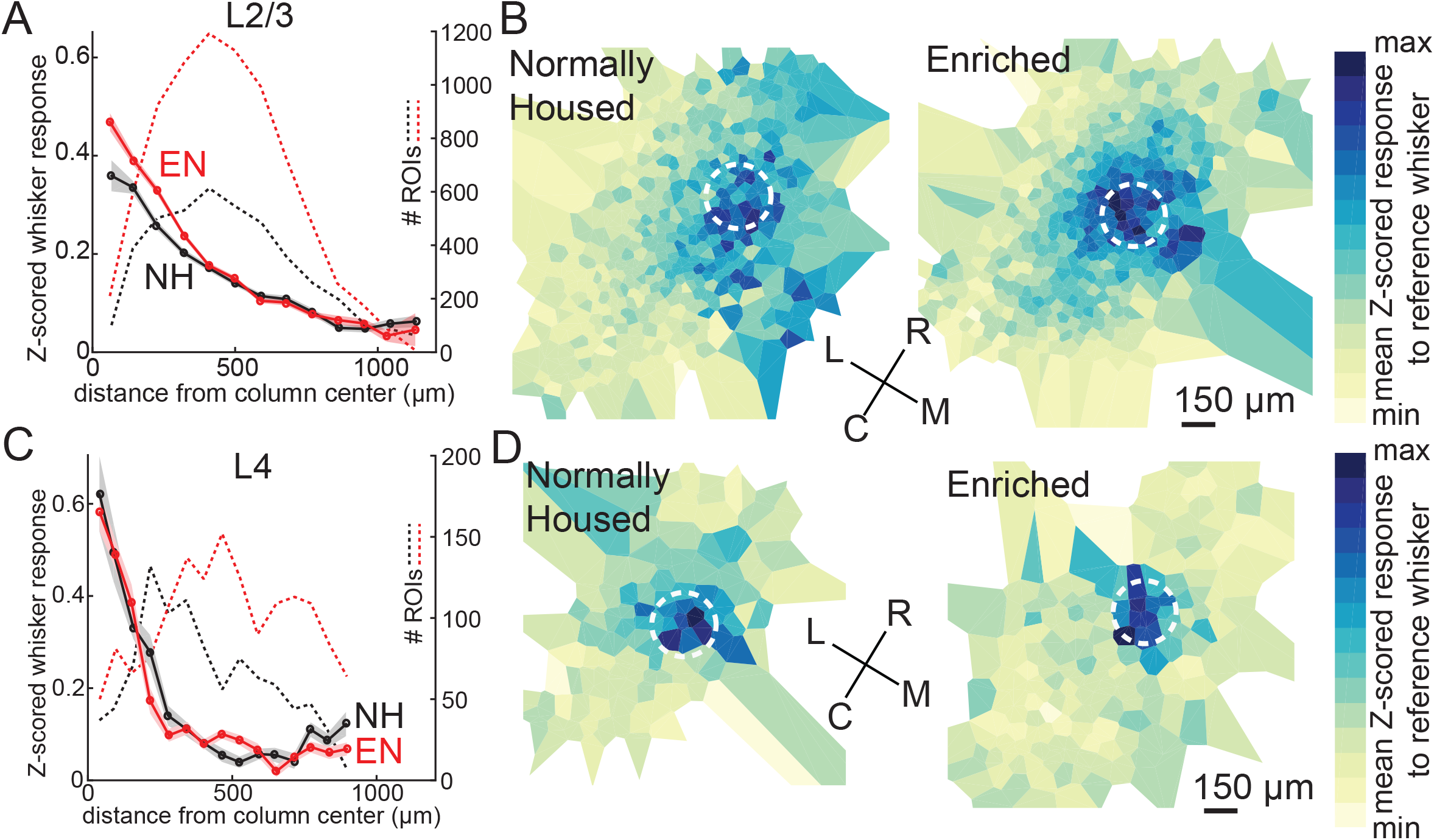
Enrichment strengthens and sharpens the whisker point representation in L2/3. **(A)** Average evoked response in L2/3 to a reference whisker (ΔF/F, z-scored to spontaneous activity in each cell) across all ROIs at increasing distances from the reference column center. ROIs are grouped into ~60 μm width bins. Shading shows SEM. Dash, number of ROIs per bin. **(B)** Mean 2D spatial distribution of evoked responses to a reference whisker for ROIs in different spatial bins around that whisker’s column. Data are compiled across all imaging fields, after alignment around the reference column center. Color shows the mean response within each bin. Dashed circle is the reference column (shown as an average barrel diameter around the column center). **(C-D)** Same as A-B, but for L4 ROIs.

In L4, mean evoked responses were stronger in the reference column center (0.62 ± 0.08 for the 0-62 μm bin), and fell off more sharply with distance than in L2/3. Slightly smaller distance bins were used, due to the spatial sampling of neurons in L4. EN did not change the peak or spatial profile of the point representation in L4 (**Fig. 5C-D**). Thus, enrichment sharpened the point representation in L2/3, but not L4, by augmenting responses in and just around the reference whisker’s home column.

### Enrichment strengthens functional column boundaries in L2/3

Prior imaging studies detected a global tuning gradient in L2/3 of S1 but no functional boundaries at anatomical column edges (Kerr et al., 2007; Sato et al., 2007). To test whether enrichment drives emergence of functional boundaries, we first examined tuning similarity (signal correlation) between pairs of simultaneously imaged L2/3 neurons. In NH mice, mean signal correlation for neuron pairs fell off modestly with intersoma distance, and was slightly higher for within-column pairs than across-column pairs. For closely spaced pairs (<150 μm apart) signal correlation was only modestly higher for within-column pairs than across-column pairs (0.652 ± 0.005vs. 0.611 ± 0.007, p<0.0001, permutation test of difference in means), reflecting the lack of functional column boundaries (**Fig. 6A**). In EN mice, mean signal correlation for within-column pairs was modestly increased (NH: 0.637 ± 0.004, EN: 0.659 ± 0.002; includes pairs at all distances; p<0.0001, permutation test), while signal correlation for across-column pairs was greatly reduced (NH: 0.541 ± 0.004, EN: 0.425 ± 0.004; p<0.0001) (**Fig. 6A, C**). Moreover, signal correlation for closely spaced cross-column pairs (<150 μm apart) was much lower than for similarly spaced within-column pairs (0.674 ± 0.003 vs. 0.529 ± 0.006, p<0.0001) (**Fig. 6A**). Thus, EN generated a tuning boundary at column edges.

**Figure 6.**
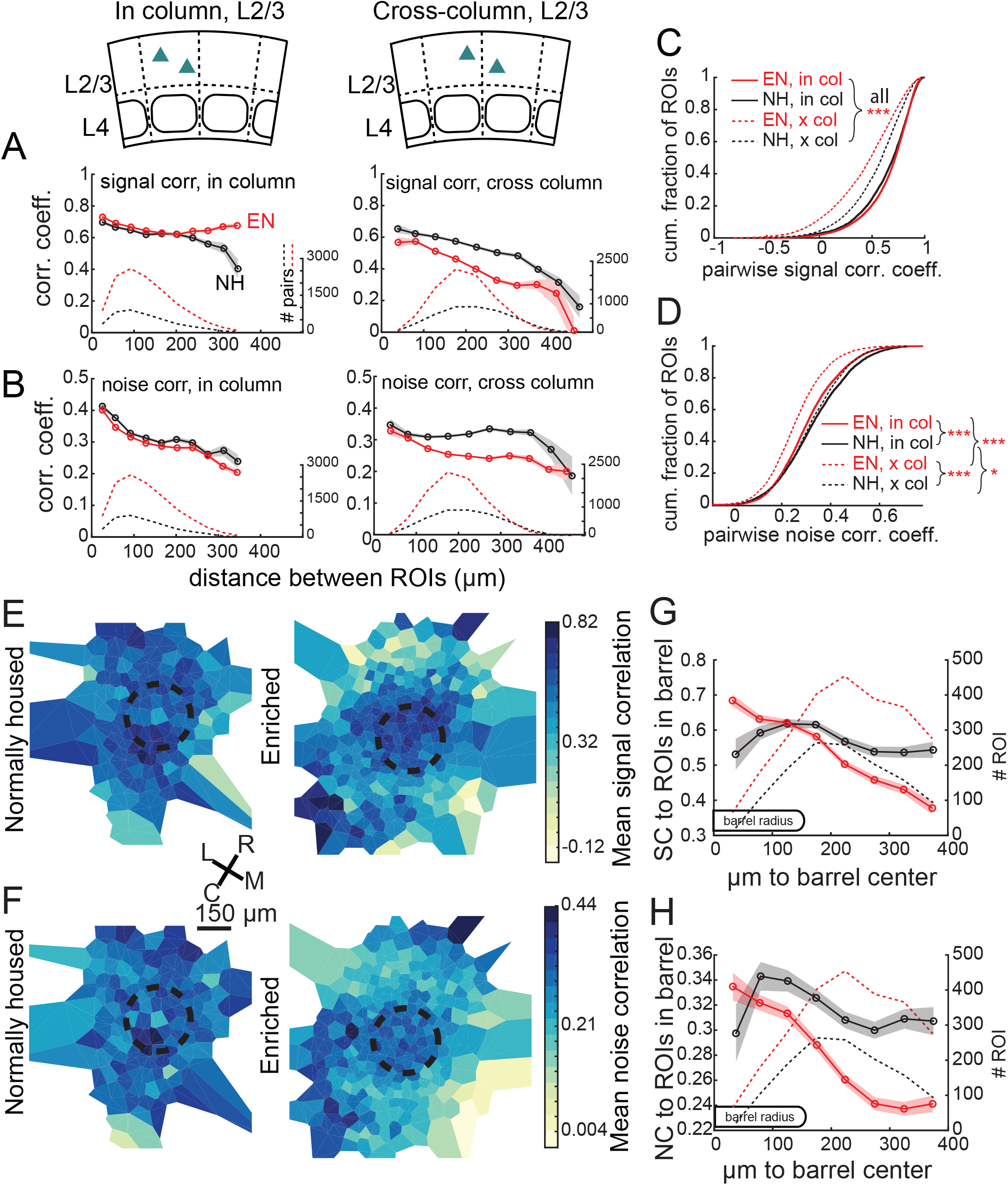
Enrichment alters the spatial structure of signal and noise correlations in L2/3. **(A)** Mean signal correlation across responsive L2/3 cell pairs as a function of inter-ROI distance, for cell pairs within a column (left) or across columns (right). Shaded regions denote SEM. Dashed, number of cell pairs in each bin. Insets show schematic of cell locations. **(B)** Mean noise correlation across all responsive L2/3 pairs, plotted as in (A). **(C-D)** Cumulative distribution of signal correlation (C) or noise correlation (D) for all within-column (in-col) and across-column (x-col) pairs. Asterisks denote difference between distributions computed by ANOVA with multiple comparisons correction. **(E)** Spatial map of mean signal correlation for sample ROIs within a position bin to all ROIs located within a reference whisker column. For each sample ROI, the mean signal correlation to all ROIs within the reference column was calculated. Sample ROIs were then clustered into spatial bins, and the mean signal correlation for each bin was plotted. The dashed circle is the reference column (shown as an average barrel diameter around the column center). **(F)** Spatial map of mean noise correlation to all ROIs within the reference whisker column, shown as in (E). **(G)** Mean signal correlation from sample ROIs to all cells in a reference column, as a function of sample ROI distance from column center. Shading is SEM. **(H)** Same as for (G), but for noise correlations. G-H show that enrichment steepens correlation gradients at column edges.

EN also generated columnar structure in noise correlations, which measure shared, stimulus-independent trial-to-trial variability between neurons, likely reflecting shared functional input (Averbeck et al., 2006; Kohn et al., 2016). In NH mice, noise correlations for L2/3 neuron pairs decreased with distance, and were similar within columns (0.334 ± 0.002) and across columns (0.327 ± 0.002). EN modestly reduced noise correlations for within-column pairs (to 0.307 ± 0.001; p<0.0001, permutation test vs. NH), but greatly reduced noise correlations for cross-column pairs (to 0.255 ± 0.001; p<0.0001, permutation test vs. NH). This effect was apparent even for cross-column pairs 100 μm apart (**Fig. 6B,D**).

To visualize functional columnar geometry based on these correlations, we calculated each neuron’s mean signal and noise correlation to the population of neurons located within the anatomical column of a reference whisker. In NH, neurons outside the reference column exhibited fairly high signal and noise correlations to the neurons inside the column. EN substantially reduced these signal and noise correlations for neurons outside the column, while preserving signal correlation and only modestly reducing noise correlation among neurons within the reference column (**Fig. 6E-F**). This generated a stronger gradient of signal- and noise-correlation in L2/3 at the reference column edge (**Fig. 6G-H**).

In L4, EN generally increased correlations between co-columnar neurons (**Fig. 7**). This was true for both signal correlations (NH: 0.681 ± 0.011; EN: 0.774 ± 0.005; p<0.0001) and noise correlations (NH: 0.241 ± 0.006; EN: 0.356 ± 0.002; p<0.0001). Too few cell pairs were imaged across columns to analyze cross-column correlations. Thus, EN drove functional differentiation of columns in L2/3, and increased coordinated activity within each column in L4.

**Figure 7.**
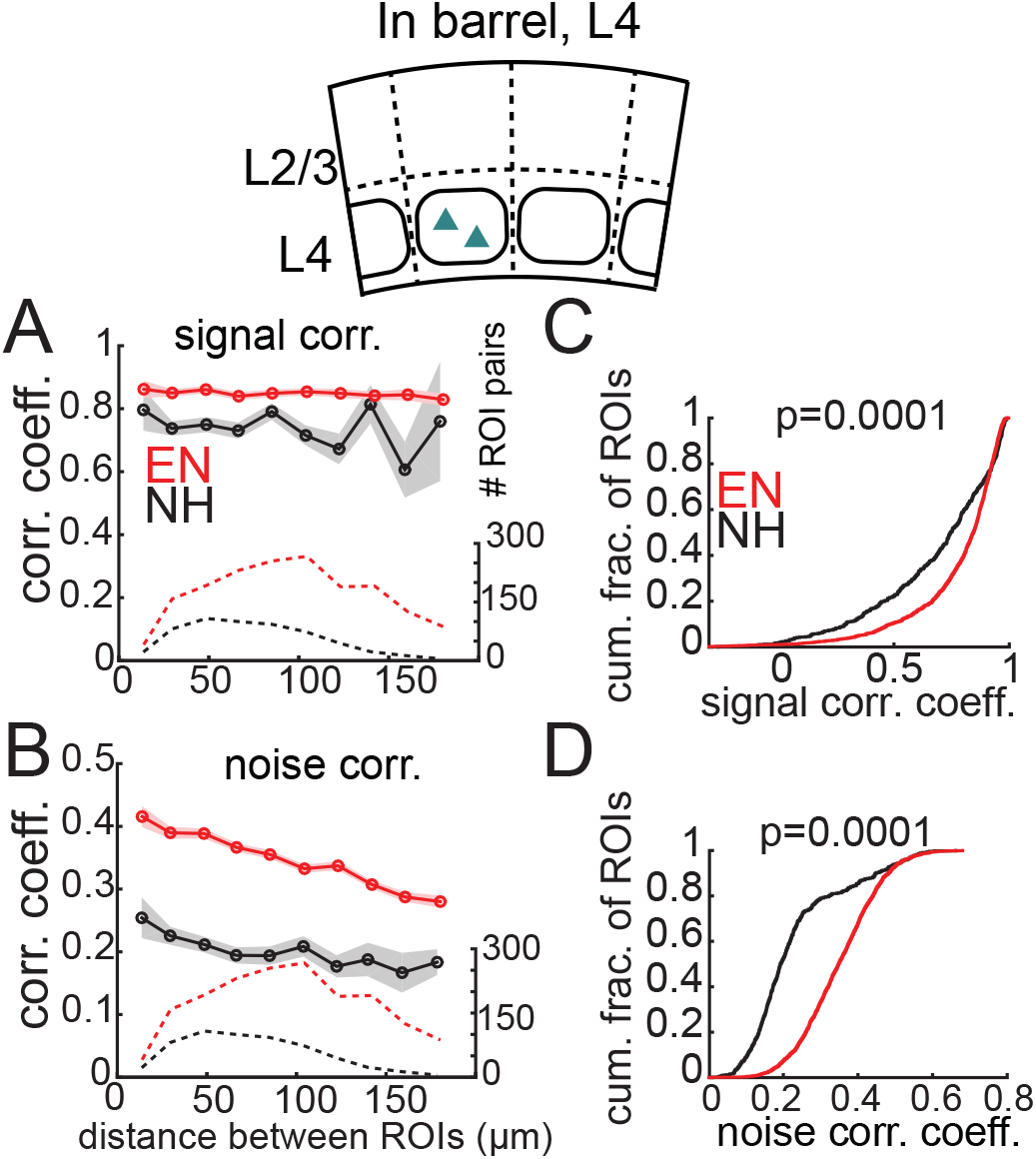
Enrichment increases signal and noise correlations within L4 barrels. **(A)** Mean signal correlation across responsive L4 cell pairs as a function of inter-ROI distance, for cell pairs within a column. Conventions as in Fig. 6A. Inset, schematic of cell locations. **(B)** Mean noise correlation across within-column pairs, plotted as in (A). **(C-D)** Cumulative distribution of signal correlations (C) or noise correlations (D) for within-barrel ROI pairs in L4.

GCaMP6s-based measures of tuning, tuning heterogeneity, and activity correlations could be contaminated by out-of-focus fluorescence from neuropil around each ROI. The extent of such neuropil contamination is difficult to measure empirically, and 2-photon imaging studies apply different methods and degrees of correction. Above, we applied no neuropil correction in L2/3, because Drd3-Cre mice had relatively low neuropil signal, reflecting GCaMP6s expression in only ~50% of L2/3 PYR cells. We did apply neuropil correction in L4 (at weight r=0.3, see Methods), because Scnn1a-Cre drove denser GCaMP6s expression, and imaging deeper in cortex required higher laser power, which could increase out-of-focus fluorescence above the imaging plane. To confirm that the major effects of the study were robust to different levels of neuropil correction, we applied neuropil subtraction in both L2/3 and L4 at 3 weights (r=0, r=0.3, r=0.7, see Methods) (**Fig. S3**). Enrichment effects on tuning heterogeneity (**Fig. 3**) and point representation (**Fig. 5**) were qualitatively unchanged across these levels of neuropil subtraction in both L2/3 and L4 (**Fig. S3A-D**). The highest level of subtraction (r=0.7) has been used in several prior studies (Chen et al., 2013) but here appeared to cause overcorrection for most ROIs in both L2/3 and L4, and drastically reduced the number of ROIs that were whisker-responsive. Effects of enrichment on signal correlations were conserved for r=0 and r=0.3, but were lost for r=0.7, perhaps because fewer cells could be analyzed (only responsive cells were analyzed). Effects of enrichment on noise correlations persisted across all neuropil subtraction levels (**Fig. S3E-F**). Thus, the major effects of enrichment were robust to methodological choices on neuropil subtraction.

### Enrichment improves population coding of columnar whisker deflections

Are the changes in mean tuning, response magnitude and activity correlations that occur in L2/3 sufficient to improve population coding of whisker input on single trials? We first asked whether EN altered the ability to detect CW deflection (relative to spontaneous activity) from single-trial ΔF/F signals from simultaneously imaged neuron populations. We built a population decoder in which each ROI was trained by logistic regression, on a subset of CW deflection and blank trials, to predict whisker deflection from ΔF/F in a 1-sec window. Performance was tested on held-out trials. Single ROIs performed above chance, with slightly better performance by EN cells than NH cells (fraction correct for NH: 0.63 ± 0.005, for EN: 0.65 ± 0.004, p=0.004, permutation test for difference in means). Population coding was tested by randomly selecting ensembles of 2-30 ROIs that were simultaneously imaged in the same field, and averaging the output of each ROI classifier to generate a population prediction on each trial. Because ROIs could be located in different columns, we averaged performance across the set of CWs for those columns. Detection performance increased with population size, and was greater for EN than NH for populations containing ≥ 3 neurons (p<0.001; permutation test of difference in means) (**Fig. 8A-C**).

**Figure 8.**
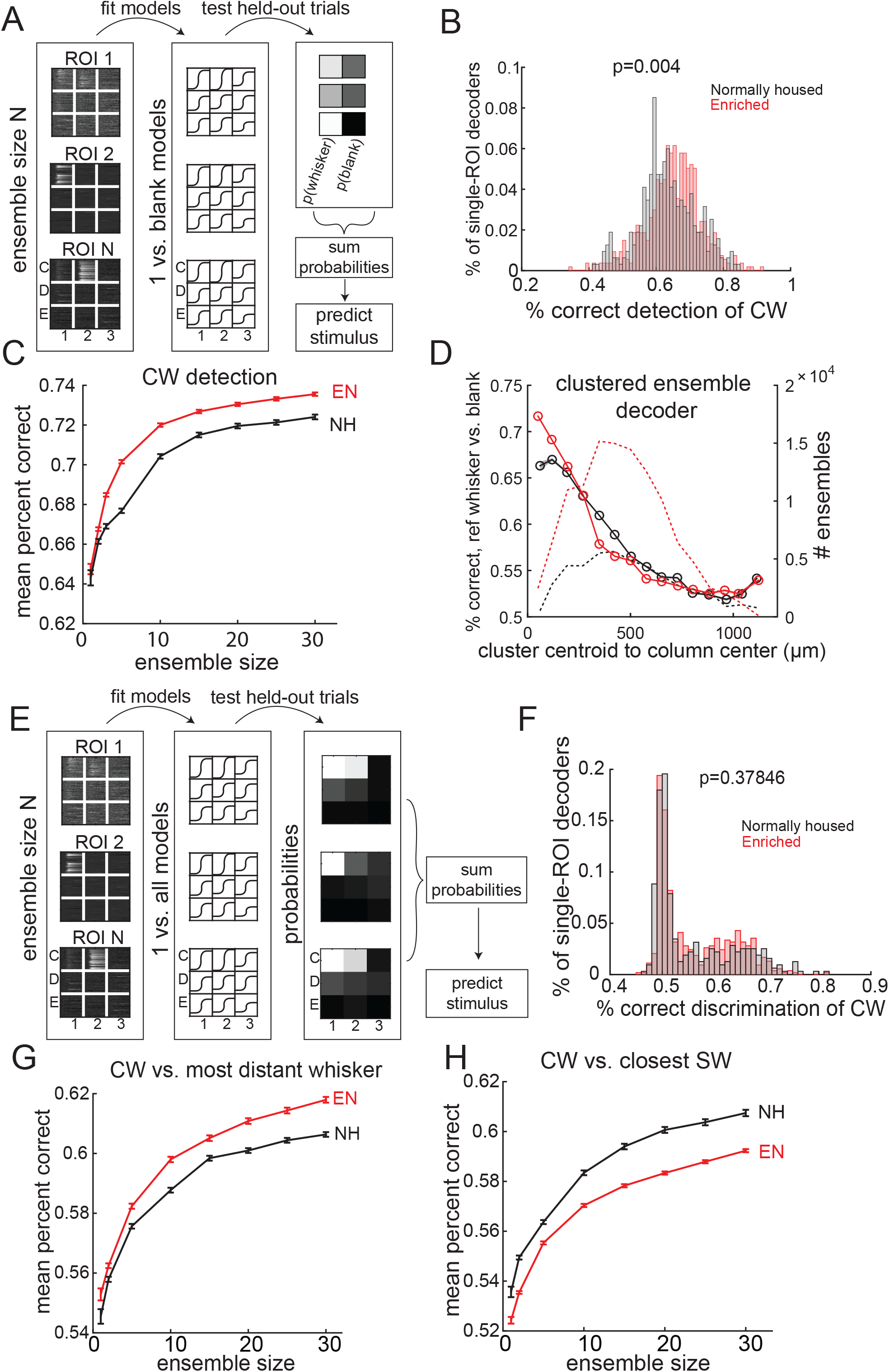
Enrichment improves population coding of CW deflections. **(A)** Design of detection decoder. **(B)** Performance of single-ROI decoders on detection of CW deflection vs. blank (no whisker deflection). **(C)** Detection performance for population decoders of varying size. Each ensemble decoder consisted of randomly positioned, simultaneously imaged ROIs, and was tested on detection of the whisker closest to the centroid of the ensemble. Bars show SEM. **(D)** Detection of a reference whisker deflection by ensemble decoders located at varying distances from the reference column. These were spatially clustered ensembles of 2-40 simultaneously imaged ROIs (see Methods). Distance bins refer to distance between reference column center and the centroid of each ensemble. **(E)** Design of discrimination decoders. **(F)** Performance of single-ROI decoders on discrimination of CW vs. any other whisker. **(G)** Discrimination performance for population decoders of varying size. Each ensemble decoder consisted of randomly positioned, simultaneously imaged ROIs, and was tested for discrimination of a reference whisker (defined as the whisker closest to the centroid of the ROIs in the ensemble) vs. the whisker corresponding to the most distant column from the decoding ensemble. **(H)** Discrimination by the same ensembles between the reference whisker and the whisker corresponding to the closest adjacent column in the decoding ensemble.

To test how detection performance varied with columnar topography, we quantified population decoding accuracy for detection of a reference whisker by spatially clustered neural ensembles (N=2-30 neurons) centered at varying distances from the reference whisker’s anatomical column. Spatially clustered, rather than randomly dispersed, ensembles were selected in each imaging field (see Methods). Mean detection performance (on average across the population sizes) was greatest for ensembles centered near the whisker column center, and fell off with distance. EN decoders outperformed NH decoders for ensembles centered at 0-77μm and 77-154 μm from a column center (p=2 * 10^−6^,2-way ANOVA), which are within an average column radius (**Fig. 8D**). But this effect disappeared and even reversed for more distant ensembles, where EN performed worse than NH (e.g., 308-385 μm bin; p=2 * 10^−6^). Thus, the strengthening and sharpening of the point representation in EN mice is sufficient to improve single-trial population detection of whisker deflections inside each whisker’s column, and causes topographic sharpening of effective decoding.

We also asked whether EN increases the accuracy of discriminating between two nearby whiskers. We built a discrimination decoder in which each ROI was modeled with 9 logistic classifiers, each of which was trained to discriminate a single whisker from the other 8 whiskers (considered as one group) based on single-trial ΔF/F. Each classifier output the probability that the corresponding whisker was deflected. Most single ROIs performed poorly at discriminating the CW vs. other whisker deflections from single-trial ΔF/F responses. EN did not improve average performance across single ROIs (NH: 0.55 ± 0.004, EN: 0.56 ± 0.003, p=0.38, permutation test of difference in means) (**Fig. 8E-F**). Population decoding was evaluated for ensembles of spatially clustered ROIs within the same imaging field. Classifier probabilities were summed across all ROIs in the ensemble, and the whisker with the highest probability was taken as the population prediction. We asked how well each ensemble discriminated each of its columnar whiskers from immediate adjacent whiskers, and from distant surround whiskers. Discrimination performance improved with ensemble size. EN increased the ability of populations to discriminate CWs from distant whiskers, but reduced their ability to discriminate CWs from adjacent whiskers (**Fig. 7G-H**). This bimodal effect on discrimination performance parallels the finding that EN strengthens multiple whisker responses at receptive field centers, but weakens receptive field edges, which sharpens the overall receptive field (**Fig. 4**). Thus, receptive field reorganization by enrichment is sufficient to predict corresponding changes in single-trial whisker discrimination.

## Discussion

### Whisker map structure in L2/3 and L4 of normally housed mice

Rodent V1, A1 and S1 exhibit dispersed, salt-and-pepper tuning organization in L2/3, without functional column boundaries (refs). We confirmed this finding in S1 of NH mice, and quantified map structure for L2/3 PYR cells, unlike prior S1 studies which did not distinguish cell types (Clancy et al., 2015; Sato et al., 2007). In NH mice, 48% of PYR cells exhibited peak tuning to the CW, and 65% responded to the CW among multiple statistically equal best whiskers (**Fig. 2**). Because of this tuning heterogeneity and the multi-whisker receptive fields of many L2/3 PYR cells (Kerr et al., 2007) (**Fig. 3**), the point representation of a whisker in L2/3 spread across multiple columns and fell off gradually without discrete columnar boundaries (**Fig. 4**). In contrast, the map among L4 excitatory cells was more spatially focused, though still contained a small degree of salt-and-pepper structure (**Figs. 2, 4**). This is consistent with clustered, topographic projections from VPM thalamus to L4, and with prior L4 spike recordings (e.g., (Armstrong-James and Fox, 1987; Simons, 1978)). Thus, whisker somatotopy is precise in L4 but more dispersed in L2/3, similar to tonotopy in rodent A1 (Winkowski and Kanold, 2013).

GCaMP6- and spike-based tuning measurements will differ because of lower single-spike sensitivity with GCaMP, and the use of train stimuli rather than single whisker deflections. In addition, some non-columnar whisker responses in GCaMP imaging may reflect slow tangential activity propagation across columns (Petersen et al., 2003) within the 1-sec response window. Despite these caveats, we found generally good correspondence between GCaMP6s- and spike-based tuning in calibration experiments (**Fig. 1F**). Imaging is essential to measure map topography among PYR cells, because dense, cell-type specific recordings with accurate localization of each soma is generally not possible using spike recordings.

### Enrichment increases receptive field and map precision in L2/3

While S1 plasticity has been well studied in response to whisker deprivation (Feldman and Brecht, 2005), the effects of enrichment are less understood. Enrichment is reported to generate larger and smoother maps of cutaneous forepaw input (Coq and Xerri, 1998), weaker but spatially sharper whisker responses (Polley et al., 2004), and larger, more overlapping whisker responses (Guic et al., 2008). These results were obtained with multi-unit recordings and intrinsic signal imaging, which lack cell type specificity and cellular-level resolution. Whether enrichment would transform the dispersed whisker map among L2/3 PYR cells into a sharper map with more distinct columnar boundaries, or a smoother map with a more gradual tuning gradient (Pluta et al., 2017), was unclear.

We found that enrichment decreased tuning heterogeneity in column centers and sharpened whisker receptive fields (**Figs. 3-4**). These changes took place both in L2/3, which is well known for robust experience-dependent plasticity, and also in L4, which retains some plasticity in adults (Landers et al., 2011; Oberlaender et al., 2012). Receptive field sharpening in L4 occurred by global weakening of all whisker responses except to the columnar whisker, while in L2/3 sharpening involved strengthening of responses to the CW and other strong whiskers, and weakening of responses to the weakest whiskers for each cell (**Fig. 4**). Macroscopically, enrichment strengthened and sharpened point representations in L2/3, by increasing whisker responses over spontaneous activity within and just around the home column (**Fig. 5**). This constitutes an increase in signal-to-noise ratio in the L2/3 map. In contrast, point representations were stable in L4. Thus enrichment sharpened the L2/3 PYR cell map and strengthened CW responses at both the microscopic (cellular-resolution) and macroscopic levels, with more subtle changes in the L4 map. These L2/3 map changes could reflect use-dependent strengthening of L4-L2/3 or local recurrent projections (Clem and Barth, 2006; Ko et al., 2013), reorganization of cross-columnar projections (Finnerty et al., 1999), or changes in POm thalamic input (Audette et al., 2018) or top-down input (LeMessurier and Feldman, 2018) to L2/3.

### Enrichment enhances functional column boundaries in L2/3

S1 contains segregated anatomical columns in L4 (Woolsey and Van der Loos, 1970)and strong radial L4-L2/3 connectivity (Bender et al., 2003; Bureau et al., 2006). Thus, it has been puzzling that prior cellular-resolution imaging (in standard-housed mice) revealed gradual somatotopic tuning gradients in L2/3 without major transitions in tuning at column edges (Clancy et al., 2015; Kerr et al., 2007; Sato et al., 2007). We hypothesized that appropriate experience could drive development of column-related functional organization in L2/3.

Functional organization at column boundaries was quantified using signal correlations, which reflect shared information coding, and noise correlations, which are thought to reflect membership in common functional networks (Averbeck et al., 2006; Kohn et al., 2016). In NH mice, signal and noise correlations fell off gradually with distance between neurons, as is typical for cortical circuits (Montijn et al., 2016; Rose et al., 2016). The spatial profile of correlations was similar within and across anatomical column boundaries, confirming gradual somatotopy and lack of sharp functional column borders (Clancy et al., 2015; Kerr et al., 2007; Sato et al., 2007). EN mice had modestly increased signal correlations within columns but substantially decreased signal correlation across columns, even for neurons located just ~100 μm apart across a column edge. Similarly, noise correlations were modestly decreased within columns, but strongly decreased across columns, even for neurons ~100 μm apart (**Fig. 6**). Thus, enrichment enhanced tuning distinctions between columns, reduced shared noise between columns, and created sharper functional boundaries at column edges, evident from both signal and noise correlations. Local tuning heterogeneity still existed in L2/3 of EN mice, but functional columnar organization was stronger.

Greater differentiation of somatotopic tuning across columns could improve the spatial resolution of whisker detection. In addition, because noise correlations are often detrimental for sensory coding (Kohn et al., 2016), their reduction may improve whisker sensory coding for downstream areas. Reduced signal and noise correlations across columns in L2/3 may reflect more rigid columnar segregation of L4-L2/3 and L2/3-L2/3 circuits, for example by reducing cross-columnar excitatory projections or enhancing cross-columnar inhibition. In L4, enrichment increased signal correlations within columns, which may reflect stronger recurrent excitation in each L4 barrel (Ashby and Isaac, 2011).

### Enrichment improves population coding of whisker deflection in L2/3

Sensory tuning and maps reflect average responses over many stimulus presentations, but perception and sensory-evoked behavioral responses occur on single trials. Single L2/3 neurons in S1 exhibit high trial-to-trial response variability (Barth and Poulet, 2012; De Kock et al., 2007), which limits coding efficiency on single trials. To test whether sharper maps predict improved neural coding on single trials, we constructed neural decoders that predicted the presence (detection) or the identity (discrimination) of a whisker stimulus from measured single-trial responses of S1 neurons. Our focus was on coding by L2/3 PYR cell ensembles, which relay information from S1 to downstream cortical areas (Chen et al., 2015).

Enrichment improved detection of CW deflections by spatially clustered PYR ensembles located in column centers (**Fig. 8**). Enrichment increased both absolute detection accuracy and coding efficiency such that in EN mice, smaller ensembles were needed for equivalent detection performance. Thus, the strengthening of CW-evoked responses, the reduction of spontaneous activity and noise correlations, and the higher proportion of CW-tuned neurons are sufficient to improve single-trial coding of whisker deflections in EN mice. Topographically, enrichment improved detection by ensembles in the home whisker column, but worsened detection by ensembles in neighboring columns (**Fig. 8D**). Thus, the columnar organization of effective decoding was sharpened, matching the sharper point representations in L2/3 of enriched S1.

Enrichment had more complex effects on single-trial whisker discrimination. Discrimination of the CW from distant surround whiskers was improved, while discrimination relative to nearby surround whiskers was not (**Fig. 8G-H**). This corresponds to the changes in mean L2/3 receptive field shape, in which enrichment decreased responses to whiskers at the flank of the tuning curve, but increased responses to the strongest central whiskers including the CW (**Fig. 4**). Because we only tested discrimination by spatially clustered (i.e., column-scale) ensembles, our results show that the tuning changes within a column are sufficient to predict altered stimulus discrimination by column-sized networks. These results do not predict how behavioral discrimination, which likely involves comparison of neural activity across many columns, may be altered by enrichment.

### Dispersed, weakly columnar maps partially reflect impoverished sensory experience

These results indicate that the dispersed, salt-and-pepper, weakly columnar somatotopy in L2/3 of S1 is partially a product of impoverished sensory experience in standard laboratory housing. Tactile enrichment strengthened and sharpened the L2/3 PYR cell map and drove emergence of functional boundaries aligned to columnar structure. Enrichment reduced, but did not eliminate, salt-and-pepper tuning heterogeneity in L2/3. Thus, salt-and-pepper organization is a robust feature of rodent L2/3 sensory maps, but its degree is flexible based on experience. Enrichment reduced both mean tuning similarity and stimulus-independent (spontaneous) firing correlations across columns. Thus, columns became more functionally independent, which predicts more accurate column-based encoding of whisker input. The circuit mechanisms for emergence of columnar structure in L2/3 remain to be identified.

## Acknowledgments

GENIE (Janelia Farm) for GCaMP6s virus, Keven Laboy-Juarez for advice on decoding analysis, Theresa Nguyen and Shilin Chen for help with histology, Paley Han for mouse colony management. This work was supported by NIH R37 NS092367.

## Author Contributions

AL and DF conceptualized and designed experiments. DF wrote Igor data acquisition software. AL conducted experiments, wrote MATLAB analysis software, and analyzed data. AL and DF wrote manuscript. DF secured funding.

## METHODS

### Animals

All procedures were approved by the UC Berkeley Animal Care and Use Committee and follow NIH guidelines. For population imaging experiments we used 17 Drd3-Cre mice (11 males and 6 females), and 11 Scnn1-Tg3-Cre mice (4 males and 7 females) (**Table 1**). Juxtacellular recordings were performed in 5 C57BL/6 mice. Mice were obtained from Jackson Laboratories.

**Table 1.** Mouse information.

|  | age at<br>imaging | sex | days<br>enriched | days<br>expressing<br>GCaMP6s | N<br>mice | N imaging<br>fields |
| --- | --- | --- | --- | --- | --- | --- |
| **Drd3-Cre; CT** | P56 ± 11 | 2F, 5M | n/a | 17 ± 2 | 7 | 10 |
| **Drd3-Cre; EN** | P63 ± 16 | 4F, 6M | 40 ± 14 | 21 ± 6 | 10 | 12 |
| **Scnn1-Tg3-Cre; CT** | P70 ± 16 | 2F, 3M | n/a | 39 ± 12 | 5 | 6 |
| **Scnn1-Tg3-Cre; EN** | P58 ± 5 | 5F, 1M | 35 ± 4 | 28 ± 5 | 6 | 8 |
| **all mice** | P62 ± 13 | 13F, 15M | 38 ± 12 | 25 ± 10 | 28 | 36 |
| **all Drd3-Cre** | P60 ± 13 | 6F, 11M | 40 ± 14 | 19 ± 5 | 17 | 22 |
| **all Scnn1-Tg3-Cre** | P64 ± 14 | 7F, 4M | 35 ± 4 | 34 ± 11 | 11 | 14 |

### Tactile enrichment

Littermates were separated into normally housed (NH) and enriched (EN) groups at weaning (P20-21). EN animals were housed with 2-3 littermates in standard mouse cages (30 x 14 x 18 cm^3^) into which a nesting enclosure and several toys were added and exchanged every 2-4 days. Toys included sticks and blocks made from wood, ridged plastic, PVC, rubber, or cement, chosen for diversity of shape and texture (**Fig. 1A**). Toys were cleaned and sterilized between uses. NH mice were housed in the same cage, with 1 littermate and no toys. All cages contained bedding and nesting material. Running wheels were not used in either NH or EN cages.

### Surgery

Before surgery, mice received dexamethasone (2 mg/kg, i.p.), meloxicam (10 mg/kg) and enrofloxican (5 mg/kg) for analgesia and to prevent infection. Mice were anesthetized with isoflurane and maintained at 37° C. The skull was cleaned and a stainless steel head-holder containing a 5-mm aperture was affixed to the skull with dental cement. Intrinsic signal imaging (ISOI) was performed through the intact skull to localize D1, D2 and D3 columns in S1 (Drew & Feldman, 2008), and a 3-mm craniotomy was made over the D2 column. A Cre-dependent GCaMP viral vector was injected at three nearby sites, at two depths each. For L2/3, we used AAV1-Syn-Flex-GCaMP6S-WPRE, injected at ~200 and ~300 μm below the dura. For L4, we used AAV9-CAG-Flex-GCaMP6s-WPRE, injected ~350 and ~450 μm below the dura. Viruses were obtained from UPenn Viral Vector Core via the Janelia GENIE Project (Chen et al., 2013). The craniotomy was sealed with a #1 glass coverslip (0.15 ± 0.02 mm thick, 3-mm diameter) cemented to the skull. Mice received post-operative buprenorphine (0.1 mg/kg, i.p.) for analgesia.

### 2-photon calcium imaging

Imaging took place 19 ± 5 days after virus injection for Drd3-Cre mice (Gong et al., 2007) and 34 ±11 days after virus injection for Scnn1-Tg3-Cre mice (Adesnik et al., 2012; Madisen et al., 2010), as required for adequate GCaMP expression in L2/3 and L4. Imaging was performed once daily observation revealed stable numbers of fluorescent neurons with few or no neurons exhibiting nuclear expression (Tian et al., 2009). 2-photon imaging was performed using a Sutter Moveable-Objective Microscope with one resonant and one galvo scan mirror. We used a 16x, 0.8 NA water-dipping objective (Nikon). Excitation was delivered with a Coherent Chameleon Ti-Sapphire pulsed laser tuned to 920 nm. Fluorescence emission was filtered with a Chroma HQ 575/50 filter and detected with a Hamamatsu photomultiplier tube (H10770PA-40). Single Z-plane images (512 x 512 pixels) were collected at 30 Hz frame rate using ScanImage (Pologruto et al., 2003).

Imaging was performed under low-dose isoflurane (<1% in oxygen) combined with the sedative chlorprothixene (0.08 mg, i.p.), with body temperature stabilized at 37°C. For L2/3 imaging, a ~400 μm^2^ field was imaged ~100-250 μm below the dura.

9 whiskers in a 3×3 array centered on D2 were inserted into 9 independent, calibrated piezoelectric actuators, positioned ~ 5 mm from the face. A plastic shield kept away nearby, non-stimulated whiskers. Whisker stimuli were presented as trains (5 impulses, 100 ms apart) of individual deflections (300 μm rostrocaudal deflection, rise/fall time 2 ms, total duration 10 ms). Trains are necessary due to sparse spiking in L2/3 (De Kock et al., 2007), (Barth and Poulet, 2012), and increase the likelihood of eliciting detectable GCaMP signals. We interleaved the 9 whisker stimuli (5 sec isi, random order) with “blank” trials in which no whisker was deflected. At each imaging field, 60 ± 10 repetitions of each whisker stimulus were presented, and imaging data was collected as approximately forty 80-second long movies. 1-2 fields were imaged in each mouse, several days apart.

After imaging was complete, the mouse was perfused with 4% paraformaldehyde, and the brain was removed, flattened, and sectioned at 50 μm parallel to the cortical surface. Sections were processed for cytochrome oxidase to reveal the barrels in L4. Imaged neurons were localized relative to barrel column boundaries by alignment using surface blood vessels and Z-stacks of GCaMP expression collected during the imaging sessions, as well as by fiducial marks at the pial surface created by high-power laser scanning after imaging was complete. We used a custom MATLAB program to localize the centroids of the 9 anatomical barrels that corresponded to the stimulated whiskers, and to calculate the X-Y coordinate of each ROI relative to these centroids.

### Cell-attached recording

For simultaneous imaging and cell attached recordings, we injected a non-Cre-dependent GCaMP6s virus (AAV1-Syn-Flex-GCaMP6S-WPRE) at ~200 and ~300 μm below the dura at one stereotaxic location in S1. After allowing ~3 weeks for expression, a 2-mm diameter craniotomy was made over S1, as described for imaging experiments. Anesthesia and recording conditions were as described for imaging. We recorded juxtacellularly from GCaMP6s-expressing L2/3 neurons under 2-photon guidance using a recording pipette (3 μm tip, 3-5 MΩ) filled with fluorescent HEPES-buffered Ringers (in mM: 126 NaCl, 20 HEPES, 2.5 KCl, 2 CaCl2, 1.3 MgSO4, 14 D(+)Glucose, 50 AlexaFluor-594, pH 7.3, 290 mOsm). A loose seal configuration was obtained, and spike-associated currents were measured in voltage-clamp mode with holding potential adjusted to maintain a holding current of 0 pA. Spikes from the loose seal recording were collected simultaneous to GCaMP imaging. Whisker stimulation was as described for imaging experiments.

### Analysis

All data analysis was conducted in MATLAB using custom-written routines unless otherwise noted.

#### Image processing and ROIs

Imaging movies were corrected for slow XY motion in Matlab using dftregistration (Guizar-Sicairos et al., 2008) (matlab file exchange). We did not observe substantial Z-axis motion. Fluorescence of each pixel was smoothed in time (moving median filter, 4 frames). Ellipsoid regions-of-interest (ROIs) were drawn manually over neuronal somata that appeared in all movies from an imaging field. The ROI signal was the mean fluorescence of its component pixels. For neuropil subtraction, a neuropil mask was created as a 10 pixel-wide ring beginning 2 pixels from the somatic ROI. Neuropil pixels that were correlated with any soma ROI (with r > 0.2) were removed from the neuropil mask. For L4 ROIs, mean fluorescence of the neuropil mask was scaled by 0.3 and subtracted from the raw somatic ROI fluorescence. For L2/3 ROIs, neuropil subtraction was not used in the main analysis, but is presented in **Fig. S3**. Fluorescence time series of all ROIs were inspected manually to remove any movies in which mean brightness decreased >~10% due to imaging errors (e.g., loss of meniscus under the objective lens). For each ROI, fluorescence time series were converted to ΔF/F defined as (F_t_/F_0_)/F_0_, in which F0 is the 20^th^ percentile of fluorescence across the entire 80-sec movie and F_t_ is the fluorescence on each frame.

#### Quantification of whisker responses, receptive fields, and response magnitude

The whisker-evoked ΔF/F signal was quantified for each ROI on each trial as (mean ΔF/F in the 1-sec period following whisker train onset) – (mean ΔF/F in the 0.5 seconds prior to the stimulus). Each cell’s 9-whisker receptive field was quantified from the median ΔF/F to each whisker across all stimulus repetitions. A ROI was considered significantly responsive to a given whisker if the distribution of evoked ΔF/F on whisker stimulus trials was significantly greater than that on blank trials. This was computed using a permutation test: for each whisker, a vector of mean ΔF/F for each stimulus iteration was combined with mean ΔF/F for each blank trial (using the same frames as for measuring evoked activity). The combined distribution of means was split into two groups, and the difference in means of the two groups was measured. This was repeated 10,000 times, and the actual difference between the stimulus and blank distributions was compared to the distribution of permuted differences in means. A difference was considered significant if it was greater than the 95^th^ percentile of the permuted distribution. P values for each of the 9 whiskers were corrected for multiple comparisons by False Discovery Rate (Benjamini and Hochberg, 1995). A ROI was considered whisker responsive if it was significantly responsive to at least one whisker above baseline activity.

For comparison of the magnitude of whisker-evoked responses across imaging fields, evoked ΔF/F was normalized to ΔF/F on blank trials. This was done by computing median evoked ΔF/F across all stimulus trials, and z-scoring to the distribution of ΔF/F on blank trials. This normalization enables comparisons across imaging fields and neurons with different levels of GCaMP expression, which yield different absolute fluorescence intensity and ΔF/F signals.

#### Normalized anatomical reference frame for spatial analysis across imaging fields

To combine imaging results across different imaging fields, ROI coordinates were transformed into a common reference frame. This was done by measuring XY position of each ROI relative to anatomical barrel centers, and rotating the coordinates for each imaging field around the column of interest to achieve the same anatomical orientation. Response or tuning measures for individual ROIs were plotted directly overlaid within this reference frame (**Fig. 3E**). Alternatively, individually ROIs were spatially binned using k-means clustering so that each bin contained the same number of ROIs, average response strength or tuning was computed for each bin, and the result was displayed as a Voronoi plot with each polygon representing a spatial bin (**Fig. 5D, Fig. 6F-G**).

#### Signal and noise correlation analysis

Signal and noise correlation were computed between all pairs of simultaneously imaged, whisker-responsive neurons that were located within (for L4 cells) or above (for L2/3 cells) any L4 barrel. Cells located within or above L4 septa were not used for correlation analysis, to avoid ambiguity in localizing the precise barrel boundary. Thus, closely spaced cross-column cell pairs were located in adjacent barrel columns, not in column and adjacent septum. To compute signal correlation, we calculated for each ROI a 9-element vector composed of its mean response to each whisker over all stimulus repetitions. Each vector was individually Z-scored. Signal correlation was defined as the Pearson’s correlation of the vectors for a pair of neurons. To compute noise correlations, we constructed for each ROI, 9 vectors (one for each whisker) containing the mean ΔF/F during the evoked window following each stimulus iteration. Each vector was individually z-scored. We computed the Pearson’s correlation for each pair of neurons separately for each whisker, then computed the mean correlation for all whiskers. Distance between neurons was calculated as the Euclidean distance (μm).

#### Neural Decoder

We constructed two neural decoders – one to detect a whisker deflection compared to spontaneous activity, and one to discriminate stimulus identity among 9 whiskers - from single-trial mean ΔF/F of individual ROIs and ensembles of ROIs. For the discrimination decoder, each ROI was represented by a one-versus-all (OVA) classifier that was trained by logistic regression to report the probability of each stimulus given the mean ΔF/F during the evoked window following a single whisker deflection, selected randomly from each stimulus iteration (structured as in (McGuire et al., 2016)). Each classifier was composed of 9 logistic functions, one for each stimulus. For each logistic function coefficients were fit by logistic regression and K-fold cross validation to relate the mean ΔF/F on each trial to the probability of given whisker having been deflected. Each iteration of model fitting was performed on 80% of trials, and performance was assessed on the remaining 20% of trials. For single ROI decoders, predictions were made by selecting the highest probability whisker on a given trial. The fitting and testing process was repeated 500 times for each single ROI decoder, and performance was averaged across iterations.

For ensemble decoding, simultaneously-imaged ROIs were clustered into ensembles either randomly or based on position within the imaging field using k-means clustering. To sample a range of ensemble sizes and positions, clustering was performed for each field while varying the number of ensembles between 2 to as many ensembles as responsive ROIs. This was repeated separately for each whisker, and positions used for clustering were normalized relative to the reference whisker. This yielded ensemble sizes of 2:42, with 4242 ± 1400 total ensembles per NH field, and 9630 ± 1501 ensembles per EN field. For ensemble decoders composed of random sets of ROIs, we trained and tested ensemble decoders of 1, 2, 3, 5, 10, 15, 20, 25, and 30 ROIs. For each ensemble size, we tested the lesser of 500 or NchooseK ensembles (where N is the total number of responsive ROIs in the field and K is the ensemble size) randomly-drawn from each imaging field. We used a maximum of 30 ROIs in order to sample as many imaging fields as possible, since the number of responsive ROIs varied across fields and several fields contained fewer than 30 responsive ROIs. To predict single-trial stimuli from ensembles, the output of each ROI classifier was normalized so that each unit had the same weight in population decoding. The population stimulus prediction was calculated by summing the probabilities of each stimulus over all units in an ensemble and selecting the stimulus with the maximal summed probability.

The detection decoder was constructed identically to the discrimination decoder, except that logistic functions were trained to discriminate one whisker from spontaneous activity, rather than one whisker vs. all other whiskers.

**Supplementary Figure 1.**
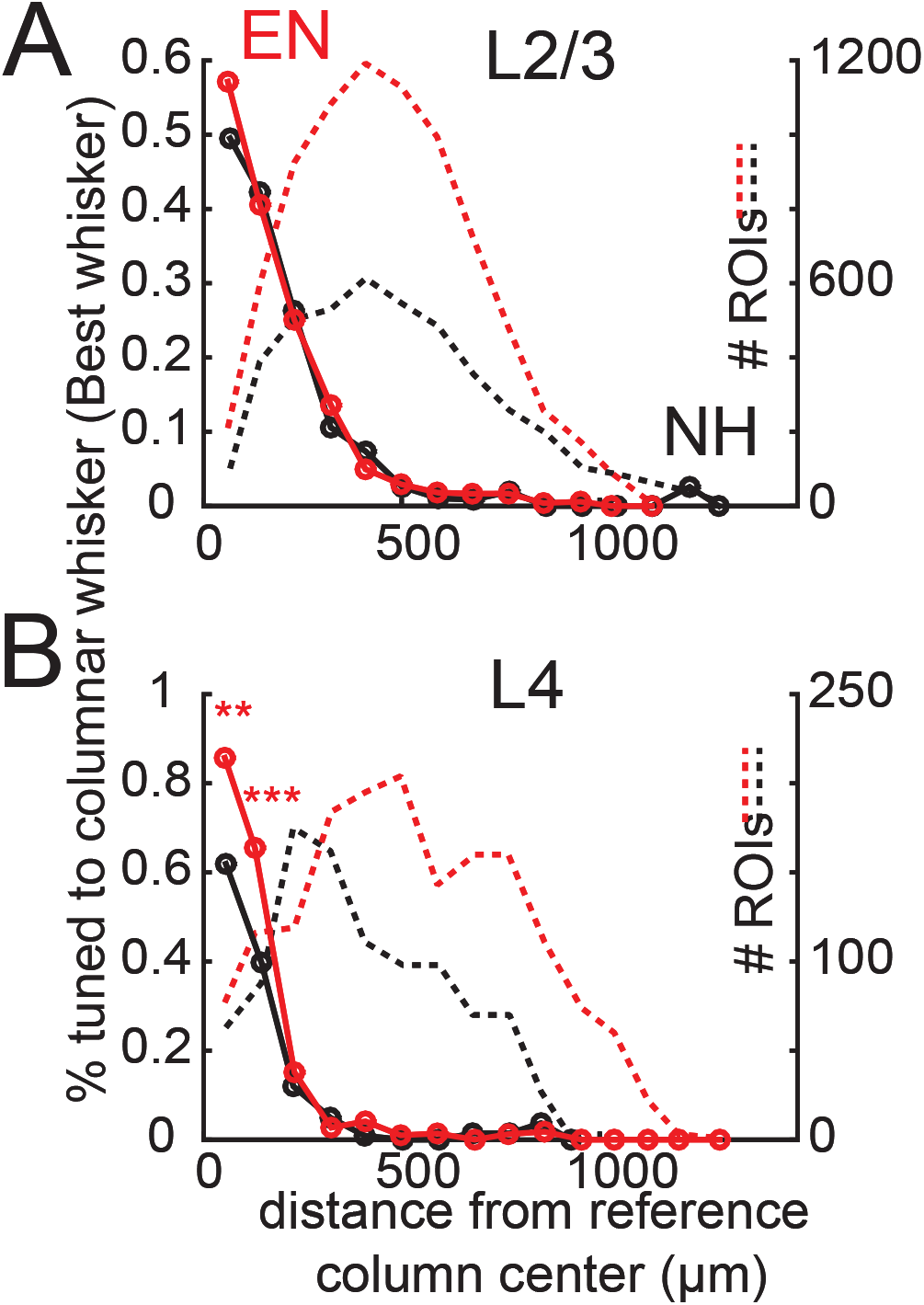
Measurements of somatotopy based on absolute best whisker. Fraction of L2/3 PYR ROIs **(A)** or L4 excitatory ROIs **(B)** that were tuned (defined as evoking the absolute largest response) to a reference whisker in EN and NH mice, binned by distance to the reference column center. Dashed lines, number of ROIs in each bin. Asterisks show significant differences between EN and NH by Fisher’ Exact Test, computed separately in each spatial bin. * p<0.05, ** p<0.01, *** p<0.001.

**Supplementary Figure 2.**
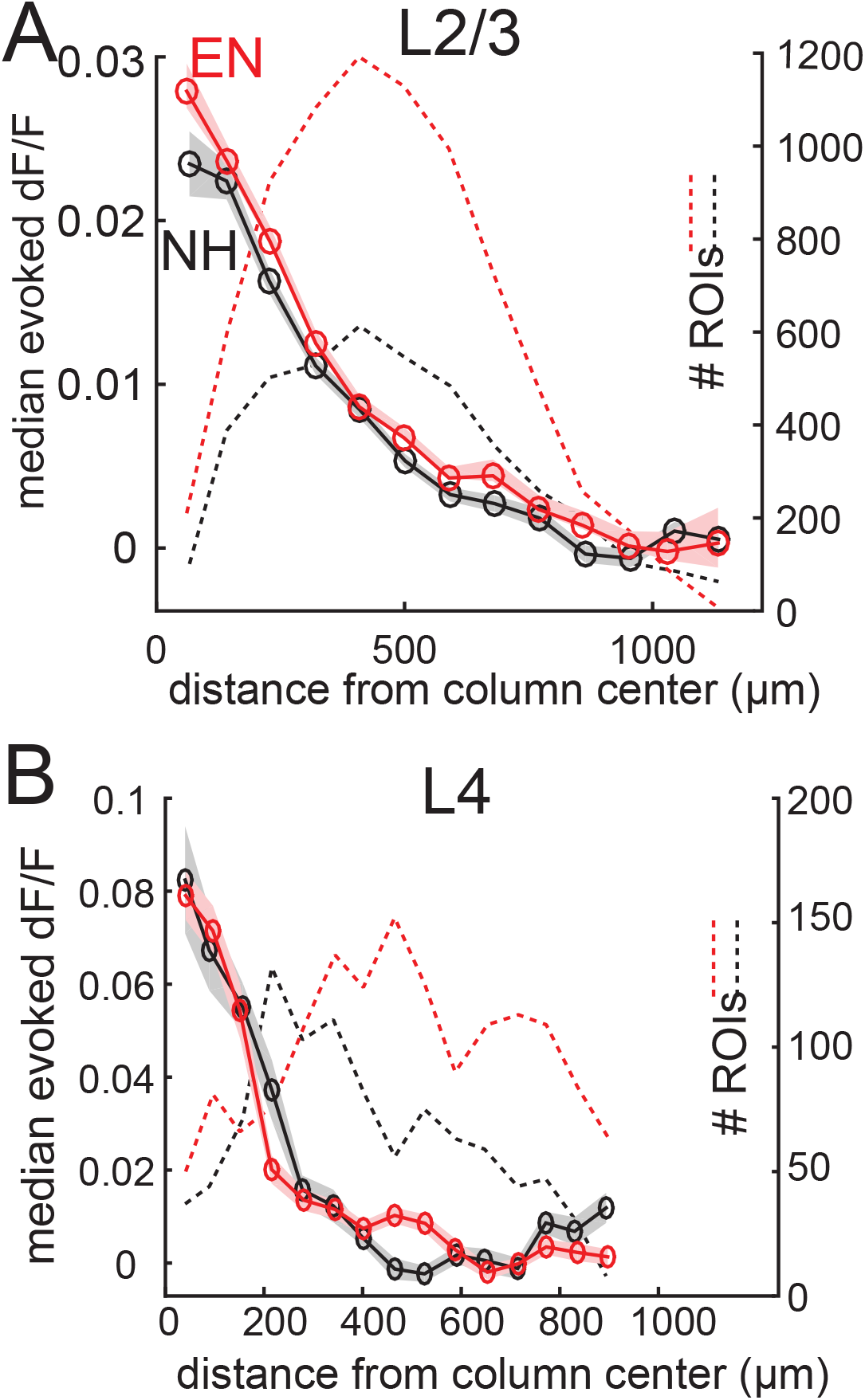
Non-normalized point representations of single whisker in L2/3 and L4. **(A)** Average evoked response in L2/3 to a reference whisker (median ΔF/F of each cell) across all ROIs at increasing distances from the reference column center. ROIs are grouped into ~30 μm width bins. Shading shows SEM. Dash, number of ROIs per bin. **(B)** As in A, for L4 data.

**Supplementary Figure 3.**
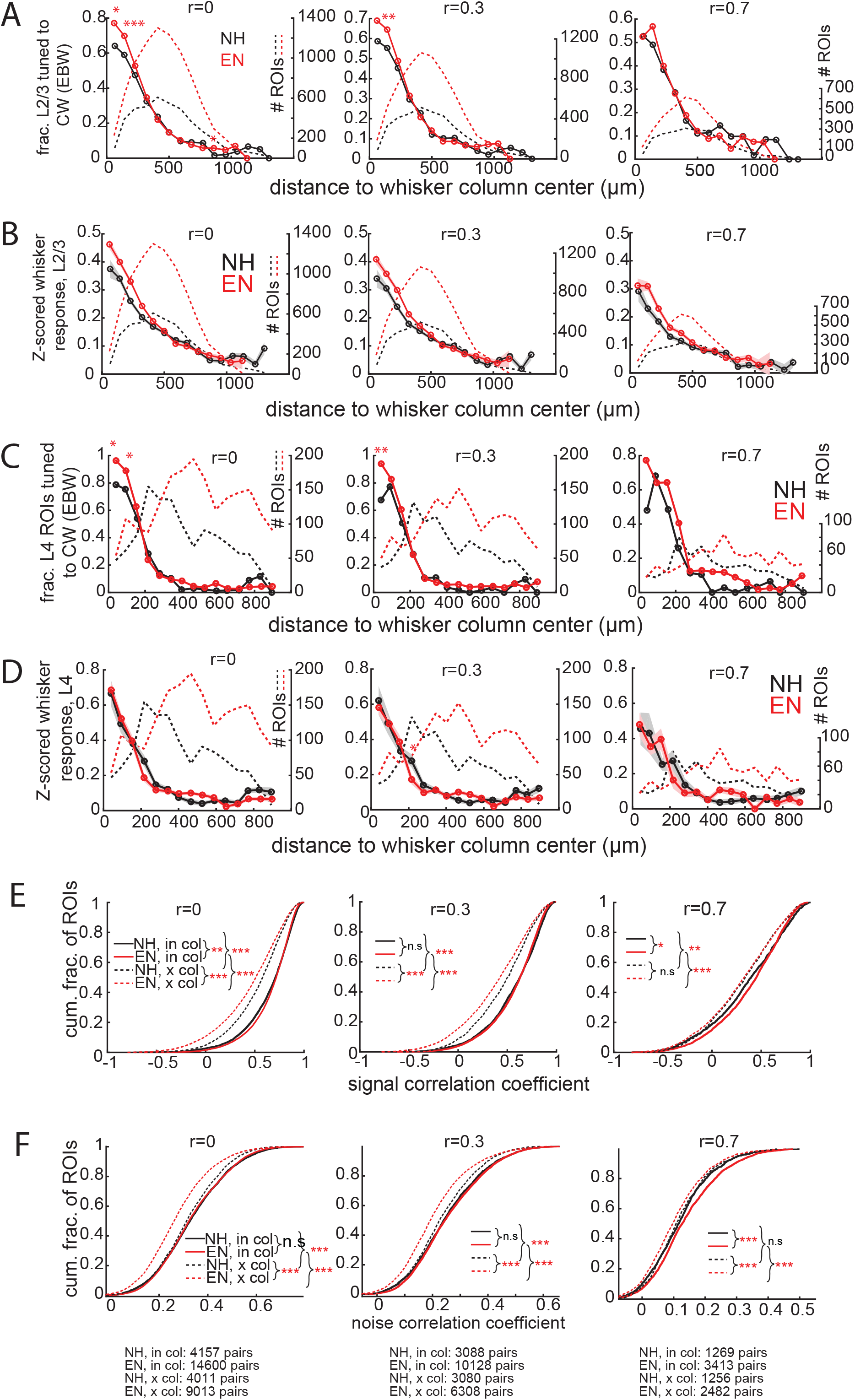
Impact of neuropil subtraction on the major effects reported in the study. **(A)** Fraction of L2/3 PYR ROIs that were equivalently tuned to a reference whisker in EN and NH mice. Left, raw fluorescence of each ROI on each frame *F(t)_ROI_* computed without neuropil signal subtraction. Middle, *F(t)_ROI_* computed according to *F(t)_ROI_* = *F(t)_ROI mask_* — r * *F(t)_NP mask_*, where neuropil signal weight r=0.3. Right, ΔF/F computed with r=0.7. Asterisks show significant differences between EN and NH by Fisher’ Exact Test, computed separately in each spatial bin. * p<0.05, ** p<0.01, *** p<0.001. **(B)** Average evoked response in L2/3 to a reference whisker (ΔF/F, z-scored to spontaneous activity in each cell) across all ROIs at increasing distances from the reference column center. ROIs are grouped into ~60 μm width bins. Shading shows SEM. Dash, number of ROIs per bin. Left, middle, and right plots are calculated with neuropil weight r=0, r=0.3, and r=0.7, respectively. **(C)** As described in (A), for L4 excitatory cells. **(D)** As described in (B), for L4 excitatory cells. **(E-F)** Cumulative distribution of signal correlation (E) or noise correlation (F) for all within-column (in-col) and across-column (x-col) pairs. Asterisks denote difference between distributions computed by with multiple comparisons correction. Left, middle, and right plots show correlation coefficients computed with neuropil subtraction weight r=0, r=0.3, and r=0.7. **(F)** Bottom row: numbers of pairs of significantly whisker-responsive ROIs in each group for cumulative distributions of NC and SC coefficients at each neuropil subtraction weight.

